# Adaptations of endolithic communities to abrupt environmental changes in a hyper-arid desert

**DOI:** 10.1101/2022.03.24.485700

**Authors:** Cesar A. Perez-Fernandez, Paul Wilburn, Alfonso Davila, Jocelyne DiRuggiero

## Abstract

The adaptation mechanisms of microbial communities to natural perturbations remain relatively unexplored, particularly in extreme environments. The extremophilic communities of halite (NaCl) nodules from the hyper-arid core of the Atacama Desert are self-sustained and represent a unique opportunity to study functional adaptations and community dynamics with changing environmental conditions. We transplanted halite nodules to different sites in the desert and investigated how their taxonomic, cellular, and biochemical changes correlated with water availability, using environmental data modeling and metagenomic analyses. Salt-in strategists, mainly represented by haloarchaea, significantly increased in relative abundance at sites characterized by extreme dryness, multiple wet/dry cycles, and colder conditions. The functional analysis of metagenome-assembled genomes (MAGs) revealed site-specific enrichments in archaeal MAGs encoding for the uptake of various compatible solutes and for glycerol utilization. These findings suggest that opportunistic salt-in strategists took over the halite communities at the driest sites. They most likely benefited from metabolites newly released in the environment by the death of microorganisms least adapted to the new conditions. The observed changes were consistent with the need to maximize cellular bioenergetics when confronted with lower water availability and higher salinity, providing valuable information on microbial community adaptations and resilience to climate change.

## INTRODUCTION

Desert microbial communities are dominated by organisms that can survive desiccation and quickly resume growth after a wetting event. The dependence on stochastic and meager inputs of liquid water exerts a marked control on how desert organisms assemble and the types of substrates they colonize. In semi-arid and arid regions, biological soil crusts (BSCs) form a thin veneer of biological activity on the top few centimeters of soils^1^. With increasing dryness, BSCs become fragmented and patchy, and microorganisms are relegated to the underside of translucent rocks such as quartz (*aka* hypolithic communities). In hyper-arid regions, hypolithic communities are replaced by endolithic communities colonizing the interior spaces of rocks^2,3^.

The hyper-arid core of the Atacama Desert in northern Chile is one of the driest regions on Earth. Primary productivity almost exclusively occurs inside halite (NaCl) nodules found on the surface of evaporitic playas (*aka* salares)^4–7^. The communities inhabiting these nodules are composed of archaea (*Halobacteria*), constituting most of the biomass, unique *Cyanobacteria*, diverse heterotrophic bacteria, and a single species of halophilic alga (*Dolichomastix*)^4–6^. The primary sources of liquid water for these communities are fog, dew, and salt deliquescence^18^. Far from representing an ecological curiosity of little significance, the Atacama halite communities are a unique example of taxonomic and functional adaptability to poly-extreme conditions of temperature, water availability, salinity, and radiation^4,8,9^.

The study of these communities can reveal new clues about the factors that control growth, taxonomic diversity, and physiological function near the dry limit of habitability. For example, the carbon isotopic composition (^13^C and ^14^C) of phospholipid fatty acids in colonized halite nodules indicates modern carbon fixation and community turnover rates of several years. However, the presence of radiocarbon enriched fatty acids also points to a significant amount of carbon recycling within the community^10^. In contrast, turnover rates in other lithic communities in hyper-arid settings, notably colder ones, are likely decades to hundreds of years or longer (e.g., Colesie, et al. ^11^, Mergelov, et al. ^12^). In addition, Archaea comprise a significant fraction of halite communities^4,13^, whereas they are mostly absent in other endolithic or hypolithic communities, and BSCs. Finally, the halite communities are dynamic and capable of responding rapidly to sudden and severe changes in environmental conditions^9^. This characteristic may explain differences in community composition within an evaporitic deposit over spatial scales of hundreds of meters to tens of kilometers^13^ and between evaporitic deposits throughout the hyper-arid core^4,14^.

Halite communities are confined to a relatively small volume within a nodule (typically 70-100 cm across and 30-50 cm thick) and, as such, each nodule represents a self-sustained microcosm that relies on limited, if any, exchange of biomass with the surrounding environment. This confinement offers a unique opportunity to relocate whole communities to different climate regions in the desert (e.g., colder, drier, higher solar irradiance) by “transplanting” entire halite nodules without disturbing the community within. Consequently, the entire community’s response to the new environment can be monitored at the community levels using molecular techniques. Few microbial ecosystems can be “forced” into new climate conditions and monitored at a different location over long periods. This paper presents the result of such a transplant experiment.

## RESULTS

Halite nodules from Salar Grande (control site) were transplanted to three other sites in the Atacama Desert, ALMA, Yungay, and Chañaral (Fig. 1, Fig. S1). We analyzed the environmental data collected at each site and the microbial communities’ taxonomic, cellular, and biochemical changes a year later (Fig. S2).

**Figure 1:**
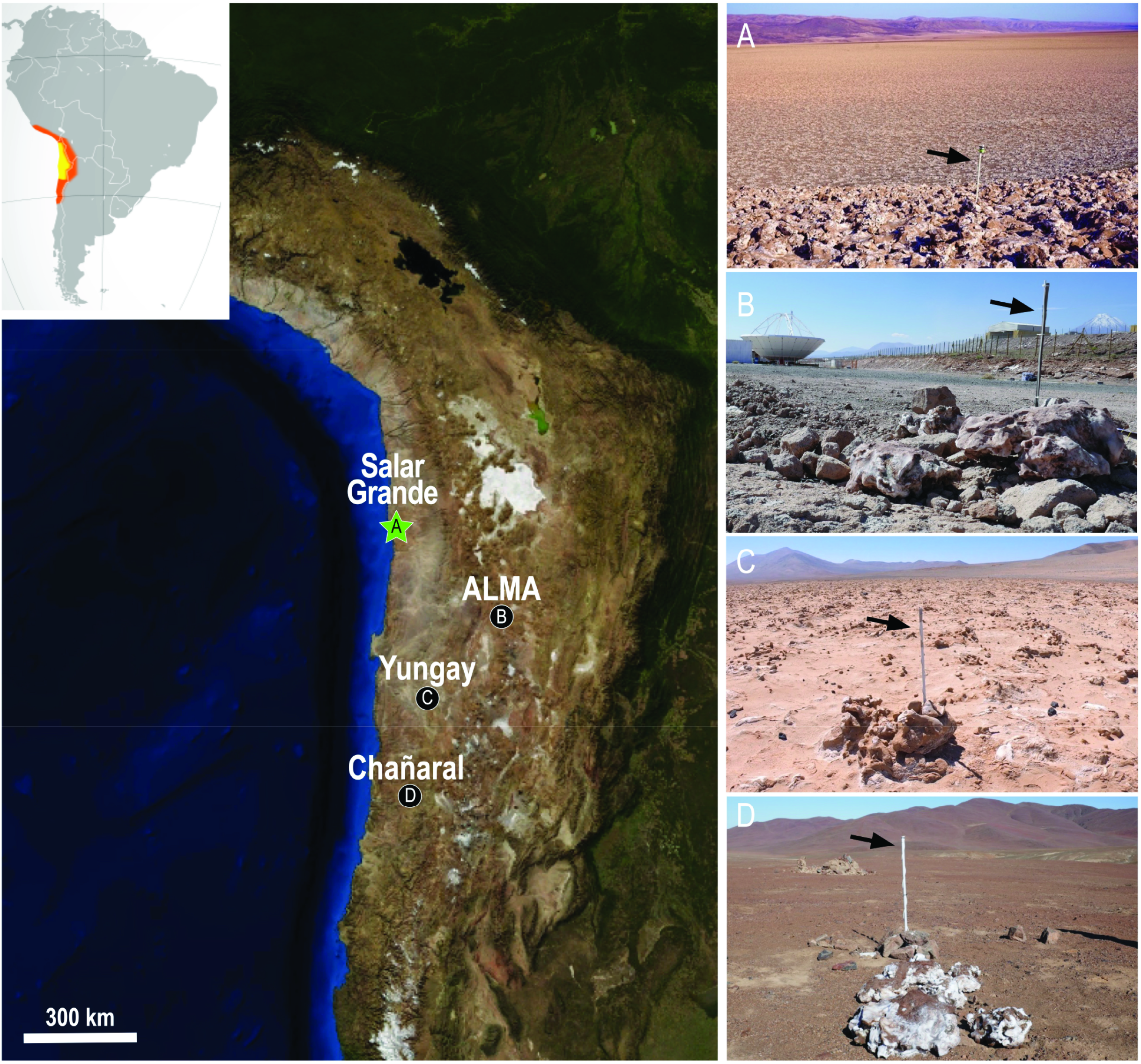
*Left*. Satellite image of the Atacama Desert in northern Chile (see insert top left) showing the location of field sites, including the control site at Salar Grande where all halite nodules originated (A), and the transplant sites at ALMA (B), Yungay (C), and Chañaral (D). *Right*. Photos of each site showing the halite nodules and the environmental sensors deployed 1 m off the ground (black arrows). Additional photos of halite nodules and the sensors installed inside are in Fig S1.

### Environmental data analysis

Different sites showed contrasting air humidity conditions RH_air_ (Fig. S3). In Salar Grande (638 masl), the maximum daily RH_air_ was above the 95% threshold for 85 days over the collection period, and the minimum daily RH_air_ values oscillated around 25%. In contrast, the ALMA site (2,885 masl) exhibited a dry regime with maximum daily RH_air_ above 95% for only 24 days and minimum daily RH_air_ values around 10%. The ALMA site also experienced three events of atmospheric precipitation (rain or snow) at approximately days 10, 160, and 280 after the transplant. During the wetting events at days 10 and 280, minimum daily temperatures were above 10°C, whereas they fell below 0°C during the wetting event at day 160. The Yungay site (953 masl) recorded the most variable T_air_ and RH_air_ readings. RH_air_ values were generally low throughout data collection, and only 16 days were recorded with RH_air_ values ≥ 95%. The Chañaral site (848 masl) was similar to Salar Grande but with more variable daily values for RH_air_ and slightly cooler daily temperatures. Chañaral recorded 239 days with peak RH_air_ ≥ 95%.

Humidity conditions inside halite nodules (RH_halite_) varied significantly between sites (Fig. S4). At Salar Grande, the mean RH_halite_ remained near 75% for the entire recorded period. No differences were observed between the top, middle, and bottom sections of the nodules. At the ALMA site, the mean RH_halite_ was initially high (75%) but dropped to values around 25% approximately 50 days after the transplant. At around day 160, corresponding to a wetting event recorded by the atmospheric sensor (RH_air_), the mean RH_halite_ suddenly increased to 75%. The RH_halite_ remained at 75% for 65 days and subsequently dropped over a period of 55 days to reach values around 25%. At approximately day 280, corresponding to another atmospheric wetting event, a second spike in the mean RH_halite_ to 75% was observed. Values remained high, albeit variable, until the end of the recorded period. Minor differences were observed between the top, middle, and bottom sections of the nodules. At the Yungay site, the mean RH_halite_ was initially high (c.a. 75%) but quickly declined to values between 50% and 60%. The mean RH_halite_ oscillated between those values for most of the recorded period, except for short intervals when it dropped to values close to 25%. Similar to ALMA, minor differences were observed between the nodules’ top, middle, and bottom sections. In the Chañaral site, the mean RH_halite_ values remained constant near 75% for the entire recorded period, which was similar to the control site in Salar Grande. Temperature conditions inside the nodules were similar at all sites and comparable to the air temperature (Fig. S4). These findings indicated that the mean RH_halite_ values point to the continuous presence of saturated NaCl brines filling the pore spaces of the halite nodules in Salar Grande and Chañaral. In ALMA, halite nodules went through three “wet periods” when mean RH_halite_ values were consistent with the presence of saturated NaCl brines (RH_halite_=75%), interspaced with two “dry periods” when mean RH_halite_ remained low. In Yungay, halite nodules progressively dried out with no observable wet/dry cycling.

Daily cumulative hours of wet and wet+light conditions further emphasized the different environmental regimes experienced by communities inside transplanted nodules compared to nodules at the control site (Fig. 2). At Salar Grande and Chañaral, conditions inside the nodules (top, middle, and bottom) were compatible with metabolic activity, both heterotrophic (wet) and photosynthetic (wet+light) for the entirety of the recorded period. In contrast, at the ALMA site, conditions inside the nodules (top, middle, and bottom) were only compatible with metabolic activity within three separate periods, each lasting 50-65 days and separated by dry spells lasting 50-100 days. At the Yungay site, conditions inside the nodules (top and middle) were compatible with metabolic activity at the beginning of the transplant experiment but slowly declined during the first 150 days. After that, only the top of the halite nodules experienced brief periods compatible with metabolic activity lasting several hours.

**Figure 2.**
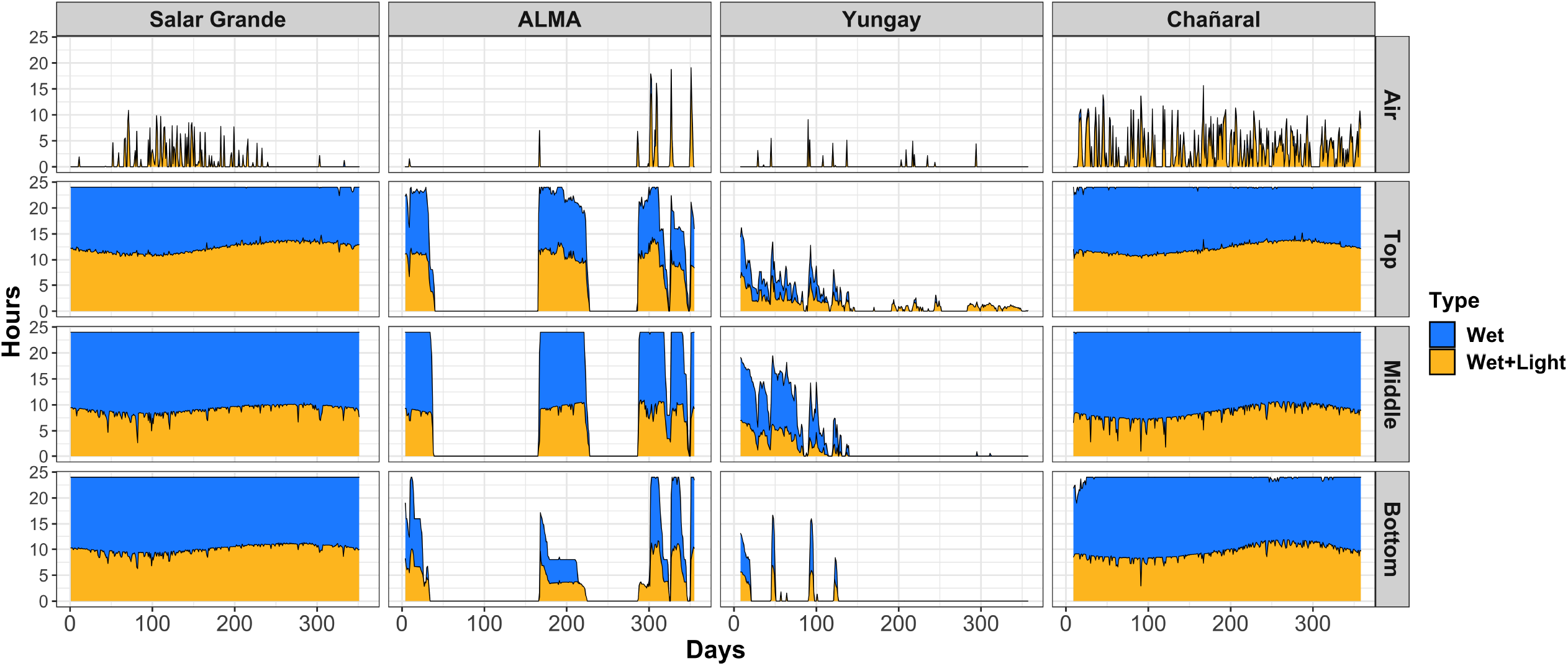
Daily hours of wet and wet+light conditions throughout data collection in the air and inside the halite nodules at each site. Salt deliquescence enables wet conditions inside halite nodules at RH>75%. In the air, wet conditions occur at RH>95%. Wet+light conditions occurred when RH values above respective thresholds co-occur with PAR>0.01.

### Long periods of extreme dryness favored *Halobacteria*

We investigated changes in the microbial community structure and function inside the transplanted halite nodules using high-throughput molecular methods. After 1 year, there was an increase in the ratio of archaea to bacteria in the ALMA transplant community and, to a lesser extent, in the Yungay transplant community, based on both Contig and MAG assemblies (Fig. 3A and Fig. 3B, respectively). In contrast, this ratio in the transplant community at Chañaral remained the same as that of the native community at Salar Grande. The higher archaeal relative abundance at ALMA resulted from an increase in the relative abundance of *Euryarchaeota* (represented mainly by *Halobacteria*) to ~80% of all contigs compared to ~60% in the Salar Grande native community (Fig. 3C; Dataset S1). In contrast, a significant decrease in *Cyanobacteria* occurred at ALMA compared to Salar Grande from ~6% to ~3% (Welch’s t-test p < 0.05, for Contigs and MAGs). *Bacteroidetes* (mostly *Salinibacter*) relative abundances were also lower at the ALMA and Yungay sites from those observed at Salar Grande and Chañaral, although not statistically significant. The same patterns were recapitulated using MAG relative abundances (Fig. 3D; Dataset S1). Overall, the variance in the data was higher in Yungay, potentially as the result of sample outliers (Fig. 3E and 3F). Ordination plots of Bray-Curtis distances showed the clustering of ALMA and Yungay transplant communities in one group and Salar Grande native and Chañaral transplant communities in another group when using contig and MAG assemblies (Fig. 3E and 3F). Although some samples such as SG2 (for contigs) and CH1 (for MAGs) appeared to be outliers, the groupings described above were statistically significant (ANOSIM of p < 0.05 for both).

**Figure 3.**
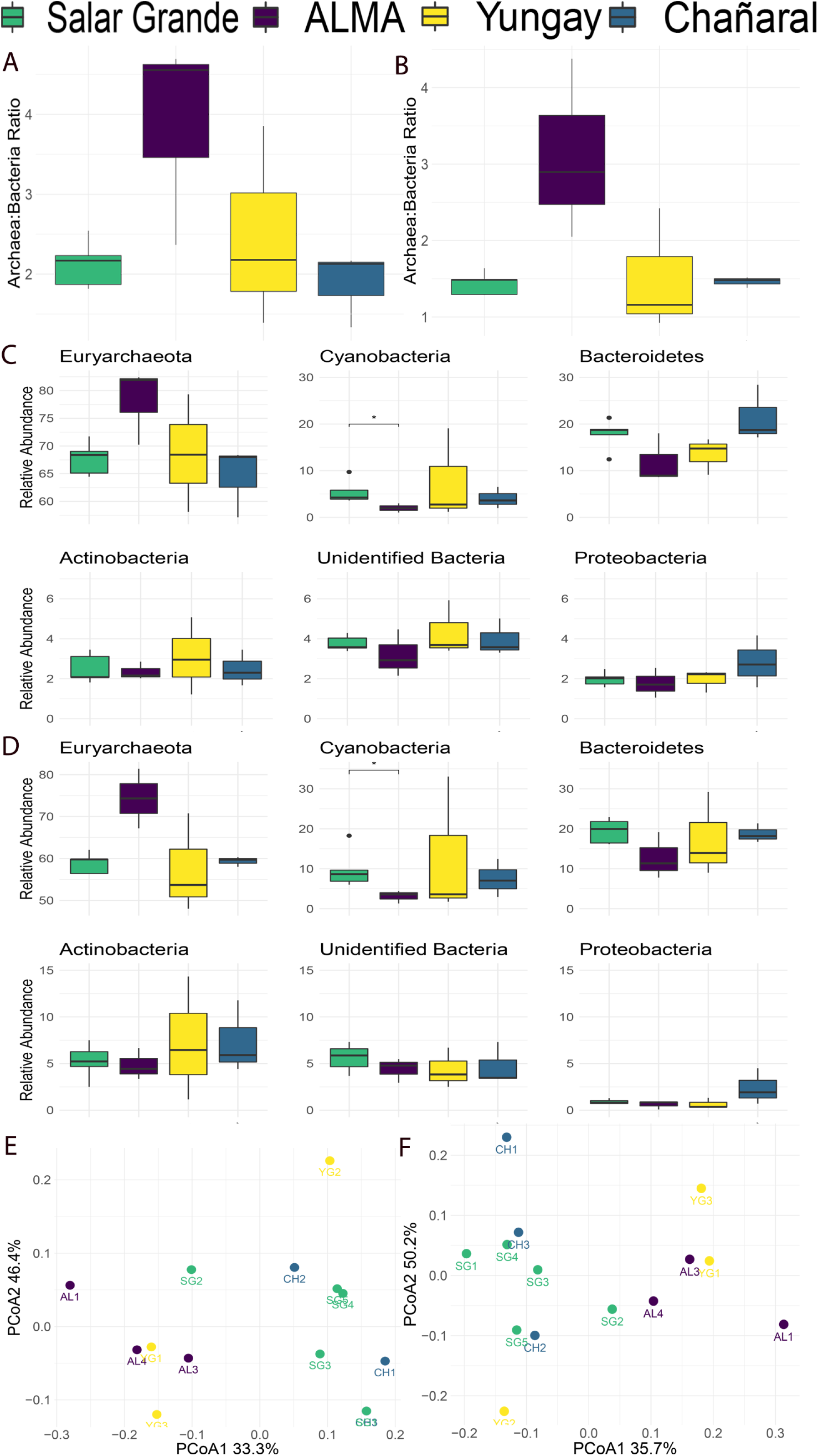
Relative abundances of halite microbial taxa across sites showing changes in community taxonomic composition after one-year exposure to different environmental conditions. Archaea: bacteria ratios in (A) contigs and (B) MAGs. Relative abundances at the phylum level in (C) all contigs and (D) MAGs. Bray-Curtis distance’s ordination plots calculated on the relative abundances of (E) contigs and (F) MAGs across sites. Error bars represent standard deviation; significance bars denote Welch’s t-test p < 0.05.

### Extreme dryness impacted the functional potential of halite communities

The functional potential of the halite communities was determined for each location using functional annotation from the KEGG Brite pathways (Fig. 4A). Overall, proteins for transport pathways such as peptide/nickel, branched amino-acids, and ABC-2 type transport systems were the most abundant in communities across all sites. Notably, the relative abundances for peptide/nickel transport systems and a HTH-type bacteriorhodopsin transcriptional activator were higher in the ALMA community when compared to other transplanted or native communities. Bacteriorhodopsin is particularly relevant to haloarchaea because it is a light-activated proton pump that can supplement the ATP budget of the cell^43^. We also found a significant decrease in the isoelectric point of the translated proteome of the ALMA community (Welch’s t-test, p < 0.05) together with an increase in potassium transport potential (*tr*K genes relative abundance) (Fig 4B and C). These adaptations are hallmarks of salt-in-strategists, which accumulate intracellular KCl to balance the high osmotic pressure of their environment, and were consistent with the observed increase in relative abundance of haloarchaea at the ALMA site.

**Figure 4.**
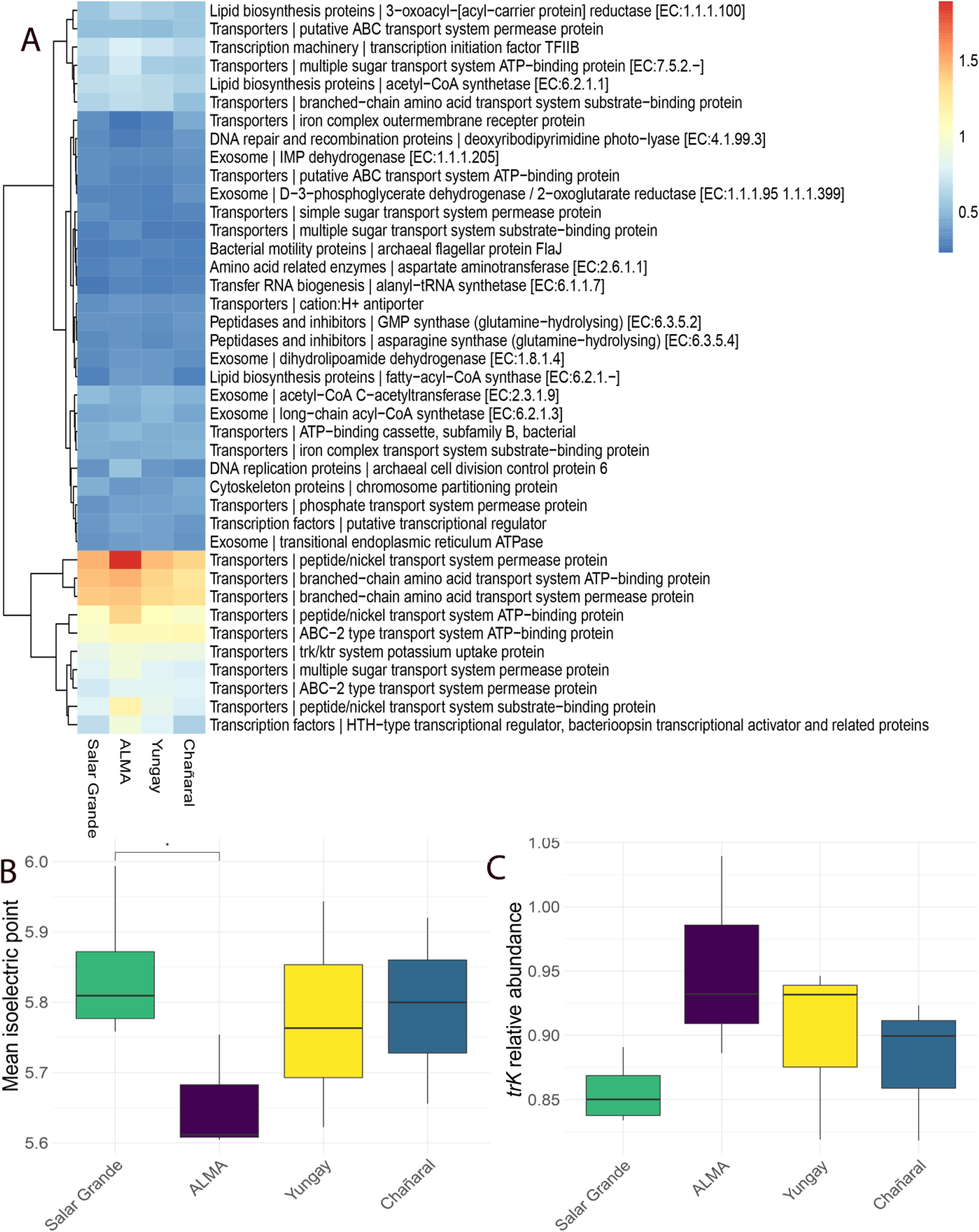
Functional changes across sites one year after transplantation of halite nodules represented by (A) relative abundances of the 40 most abundant KEGG Brite pathways in all contigs, (B) weighted average of predicted protein isoelectric points of proteins encoded in all contigs, and (C) potential potassium uptake inferred from *trK* genes relative abundance quantified in all contigs.

### Few specialized MAGs dominated halite communities

One hundred and four good quality MAGs (completeness ≥ 70% and contamination ≤ 5%) were recovered from the halite community metagenomes after binning, quality filtering, and dereplication (Dataset S2). Taxonomy annotation at the phylum level revealed that most MAGs belonged to the *Euryarchaeota* (69/104 MAGs). The changes in relative abundance of most of the MAGs were site-specific (Fig. S5). We also found six MAGs that were highly abundant in at least one site and showed the highest changes in relative abundance between the control site (SG) and the transplanted sites. We called them “specialized MAGs”; MAG.i.4, MAG.i.34, and MAG.i.36 were annotated as *Euryarchaeota*, MAG.i.2, and MAG.i.39 were annotated as *Bacteroidetes*, and MAG.i.27 were annotated as *Cyanobacteria* (Fig. 5A).

**Figure 5.**
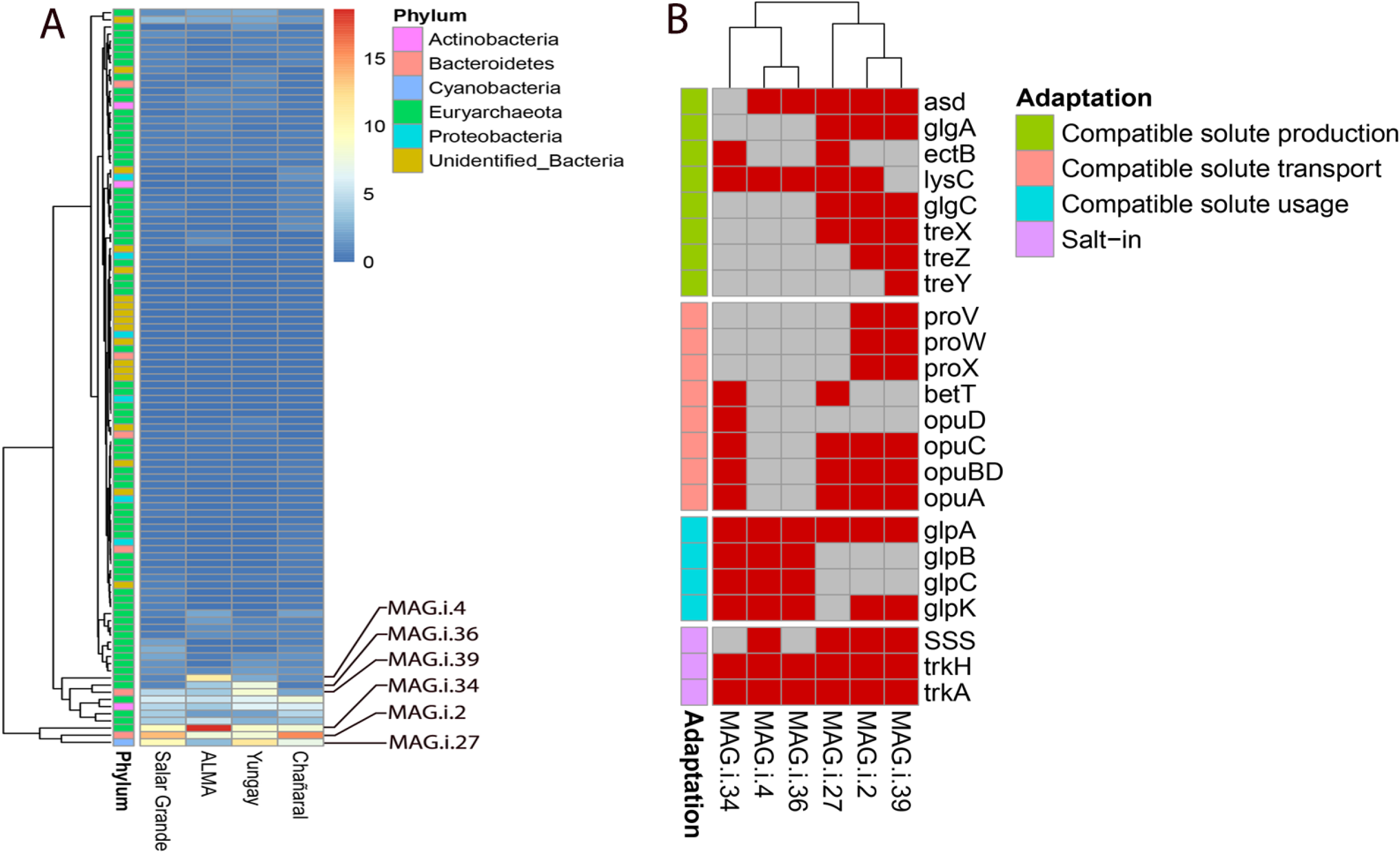
Taxonomic and functional characterization of MAGs across sites. (A) Taxonomic annotation and relative abundance of MAGs across sites; labeled are the most abundant site-specific MAGs. (B) Presence and absence of genes involved in compatible solute production, transport, and utilization in the site-specific MAGs.

Deeper taxonomic identification of the specialized MAGs with the GTDB-tk (ANI > 98% with the closest relative MAG in the GTDB; Table S1) showed that the archaeal MAGs were exclusively represented by *Halobacteriales* and included *Natronoarchaeum* (MAG.i.4), *Halococcus* (MAG.i.34), and *Halorussus* (MAG.i.36) genera (Fig. S6A). The *Bacteroidetes* MAGs belonged to the *Salinibacteraceae* family and were clustered with unidentified genomes related to *Salinibacter* (MAG.i.2) and *Salisaeta/Logimonas* (MAG.i.39) genera (Fig. S6B). The cyanobacterial MAGs (MAG.i.27) belonged to the *Rubidibacter* genus and were closely related to the *Halothece* and *Euhalothece* genera (Fig. S6C). The increased relative abundance of the specialized MAGs was site-specific. For example, *Halococcus* and *Natronoarchaeum* MAGs relative abundance was ~18% and ~11% at ALMA, respectively, compared to ~9% and ~0.9%, respectively, in the native nodules of Salar Grande and the transplants of Yungay and Chañaral (Fig. 5A). The relative abundance of *Halorussus* MAGs also increased from ~0.7% in Salar Grande native nodules to ~4% in ALMA and ~7% in Yungay transplants. There were also MAGs that decreased in relative abundance after the transplant. *Salinibacter* MAGs decreased from ~13% in Salar Grande native nodules to ~8% in the ALMA and Yungay transplants, and *Rubidibacter* MAGs declined from ~10% in Salar Grande native nodules to ~3% in ALMA transplants (Fig. 5A).

Protein encoding genes for osmoregulation were identified in the specialized MAGs with distribution indicating different osmoadaptation strategies. All the archaeal MAGs (MAG.i.4, MAG.i.34, and MAG.i.36) encoded ion transport genes, including for a sodium: solute symporter of the SSS family and for the Trk potassium transport system (*trkH* and *trkA*), both of which are functional traits of salt-in strategists. These MAGs also encoded genes for a glycerol kinase (GlpK) that catabolizes glycerol to glycerol-3-phosphate (G3H) and a G3P dehydrogenase (GlpA, GlpB, GlpC), potentially conferring the ability to use glycerol as a carbon source^44^. *Halococcus* (MAG.i.34), the most abundant MAG in ALMA transplants, also encoded genes for the uptake of several compatible solutes; these included genes for a OpuC transporter for a broad spectrum of compatibles solutes including choline, ectoine, glycine-betaine, OpuA and OpuD transporters for glycine-betaine, opuBD for choline, and betT for choline, glycine-betaine, and proline^45^ (Fig. 5B). In contrast, these genes were not found in the *Natronoarchaeum* (MAG.i.4) and *Halorussus* (MAG.i.36). MAGs from the *Salinibacteraceae* (MAG.i.2 and MAG.i.39), also salt-in strategists, encoded genes for a sodium: solute symporter and the Trk potassium transport system. The *Rubidibacter* MAGs (MAG.i.27) encoded for genes for trehalose biosynthesis such as *glgA* (starch synthase), *glgC* (glucose-1-phosphate adenylyltransferase), and *treX* (isoamylase)^46^; and the ectoine biosynthesis such as *asd* (aspartate-semialdehyde dehydrogenase), *ectB* (diaminobutyrate-2-oxoglutarate transaminase), and *lysC* (aspartate kinase)^47^.

### *Dolichomastix* and *Halovirus* relative abundances across sites

Algal contigs, assembled from the halite metagenomes, belong to a unique alga, *Dolichomastix,* whose MAG was previously reported from Salar Grande^15^. We quantified *Dolichomastix* relative abundance at each site by mapping sequence reads to the genome sequence assembled in 2021. There was no statistical difference in the relative abundance of *Dolichomastix* across sites; however, its mean relative abundance was lower in the ALMA and Yungay communities (~13%) than those of Salar Grande and Chañaral (14.5%) (Fig. 6A).

**Figure 6.**
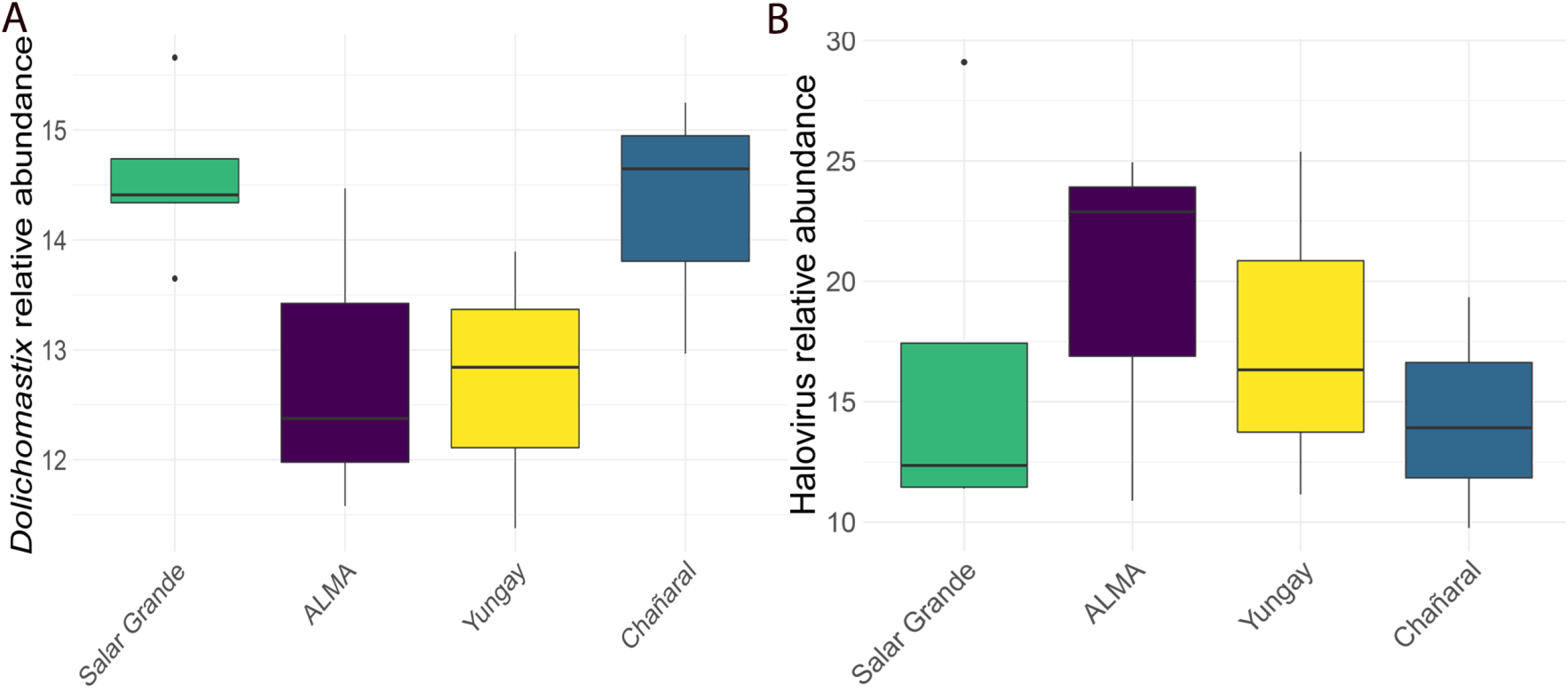
Algal and viral contigs quantification. (A) Relative abundance of *Dolichomastix* genomes in the halite communities, and (B) *Halovirus* contigs relative abundance across sites. Standard deviation is represented by error bars and the significance tested by Welch’s t-test.

Using virsorter2, we found 571 viral contigs across all samples. Of these, 101 were annotated as *Halovirus* with Cenote-Taker2, 119 were unknown viruses, and 334 were unknown phages. The relative abundance of all viral contigs was similar across all sites (Dataset S3); however, the relative abundance of *Halovirus*, the most abundant known virus family in the halite communities, varied across sites. There was an increase in *Halovirus* relative abundance in ALMA and Yungay that matched the increase in the relative abundance of haloarchaea at those sites (Fig. 6B).

## DISCUSSION

Transplanting halite nodules to different sites of the Atacama Desert triggered an adaptive response of the halite community, leading to changes in taxonomy, cellular and biochemical properties related to local environmental conditions. All the halite nodules transplanted from Salar Grande contained liquid brines in their pore spaces, as evidenced by the high RH_halite_ values at the start of the experiment. After the transplants, dry conditions and specific wetting events resulted in three distinct periods of water availability inside the nodules at ALMA. In contrast, halites at Yungay showed a progressive loss of water available to microorganisms, followed by a protracted period of desiccation. Halite nodules at Salar Grande and Chañaral maintained their water content for the entire duration of the experiment.

Water dynamics inside the nodules had an apparent effect on the composition and functional adaptations of the halite communities, with archaea and bacteria showing different trends. *Euryarchaeota*, mainly *Halobacteria*, the dominant taxon at Salar Grande, increased significantly in abundance at the sites where moisture content decreased over time (ALMA and Yungay), while it remained unchanged at Chañaral, where moisture content stayed the same. However, reduced water availability did not affect all *Halobacteria* in the same way; *Halococcus, Halorussus*, and *Natronoarchaeum* were the main taxa that increased in relative abundance at the driest sites. On the other hand, *Bacteroidetes*, mainly represented by *Salinibacter*, was the dominant bacterial group at Salar Grande and showed a slight decline at the drier sites. A reduction in *Bacteroidetes* abundance was also noted by Robinson, et al. ^4^ when comparing halite communities from Salar Grande with native halite communities at the drier Yungay site. On the other hand, Uritskiy, et al. ^9^ reported an increase in the relative abundance of *Bacteroidetes* in halite nodules at Salar Grande with increasing water availability after a rainfall. The observed trends recapitulate the changes in community structure observed elsewhere along salinity gradients, with haloarchaea dominating at the highest salinities (lower water content), while the relative abundance of *Bacteroidetes* remains practically unchanged^48,49^.

Cyanobacteria were also affected by changes in water content, with a decline in relative abundance at the ALMA site but an increase at the Yungay site. The latter is consistent with Robinson, et al. ^4^, showing that *Cyanobacteria* were more abundant in native halite nodules at the Yungay site than at the Salar Grande site and supports the notion that these photosynthetic microorganisms are generally well adapted to extreme dryness^13^. Their decline at the ALMA site appears counter to that argument but may be explained by the higher energy demand imposed by wet/dry cycling (see below) compounded by the colder conditions at this site. We also found a decrease in the algae relative abundance at the driest sites indicating a potential dry limit for eukaryotes, as previously reported^4^.

Wet/dry cycling is a significant stressor for all organisms, even for desert organisms specialized to survive extreme dryness^50,51^. Inside halite nodules, wet/dry cycling also causes important salinity and osmotic pressure changes. Changes in salt concentration are energetically expensive. Consequently, the amount of energy generated during dissimilatory metabolism and the mode of osmotic adaptation are strong selective factors in environments with high (and fluctuating) salt content^52^. The observed increase in the relative abundance of salt-in strategists, although primarily members of the archaea, in transplants at the ALMA site, and, to a lesser extent, at the Yungay site are consistent with the need to maximize cellular bioenergetics when confronted with changes in water availability. Indeed, the salt-in strategy has a lower energy cost than other adaptations to high osmotic pressure, such as salts extrusion and compatible solutes, which requires the active synthesis or uptake of compounds such as sugars, polyols, and amino acids^52^. The advantage of salting-in should be more decisive in environments where organisms go through multiple wet/dry cycles over a relatively short time, like at the ALMA site, since such fluctuations are energetically taxing. But the use of compatible solutes might also be effective in situations where organisms progressively dry out and then remain desiccated, as was observed at the Yungay site because organisms have ample time and energy to synthesize compatible solutes. This latter point would explain the observed decline of *Cyanobacteria* at the ALMA site (wet/dry cycling; higher energy cost) and their relative increase at the Yungay site (progressive desiccation; lower energy cost). Despite being salt-in strategists, *Bacteriodetes* decreased in abundance at the driest sites, which is in line with reports that correlated increases in *Euryarchaeota* relative abundance with *Bacteroidetes* declines in the relative abundance at higher salt concentrations^53–55^. The vast increase in abundance of haloarchaea at the driest sites most likely outcompeted the adaptation potential of a few genera or families of *Bacteriodetes* salt-in strategists.

The increasing dominance of salt-in strategists at the ALMA site was also reflected in the functional potential of the halite communities. Salt-in strategists uptake KCl to balance the high osmotic pressure of their environment^43,56^, and *trK* transport systems are essential in obtaining the high cytoplasmic concentration of K found in these microorganisms^51,57^. Consequently, we saw an increase in the relative abundance of *trK* genes in the ALMA and Yungay communities, with a more significant increase at ALMA consistent with the succession of wet/dry cycles compared to the gradual decline in water content observed at Yungay. A function related to energy production that was found enriched in the ALMA community transplants was the *HTH* transcriptional activator of bacteriorhodopsin. Bacteriorhodopsin is an essential protein complex in haloarchaea that acts as a light-dependent proton pump to supplement the energy budget of haloarchaea; it illustrates another adaptive strategy to meet the energetic demands at high and fluctuating salt concentrations^43^. The adaptive advantage of salt-in strategists was also reflected in the lower isoelectric point values measured in the meta-proteome of ALMA transplants (mean pI ~ 5.4 in comparison of ~5.8 at other locations). The proteins of salt-in strategists proteins are enriched in acidic amino acid residues at their surface compared to their non-halophilic homologs to maintain stability and activity at high salt concentrations^9,58,59^. The same dynamic was reported in other extreme environments where the dominance of salt-in strategists, *Euryarchaeota* and *Salinibacter,* was correlated with lower isoelectric point values of the community’s metaproteome^9,55,60^.

Poor nutrient availability is another constraint for communities that rely on the primary production of *Cyanobacteria* and algae, as is the case of in halite nodules^5,6,13^. Haloarchaea mainly use amino acids as carbon sources^61^, transported inside the cell by ABC-type transporter systems. ABC-type transporters were the most abundant pathways across all halite communities, and those related to peptide/nickel transport increased in response to water loss in the ALMA and Yungay transplants. Additionally, our MAG functional analysis revealed that three dominant archaea at the driest sites (*Halococcus, Halorussus*, and *Natronoarchaeum*) encoded for genes for a glycerol-3-phosphate dehydrogenase (*glpA, glpB, glpC*) and a glycerol kinase (*glpK*), all essential proteins for the use glycerol as carbon source^39,62^. *Halococcus*, the most abundant MAG in the ALMA transplant, also encoded transporter genes for various compatible solutes (*opuA, opuBD*, *opuC*, and *opuD*), indicating their potential use as additional carbon sources by this organism.

These findings support the conclusion that opportunistic salt-in strategists took over the halite communities at the driest sites. They most likely benefited from carbon sources, such as glycerol, choline, ectoine, glycine-betaine, and proline, newly released in the environments by the death of microorganisms the least adapted to desiccation or wet/dry cycling and colder conditions recorded at Yungay and ALMA, respectively. These opportunist salt-in strategists were all members of the archaea and were site-specific, harboring the functional potentials best adapted to the new environmental conditions following the transplant. The decrease in relative abundance of cyanobacteria, algae, and *Bacteriodetes* we observed at the driest sites supports this idea. Drastic changes in halite communities were previously reported, albeit in the opposite direction, resulting from a massive rain event^9^. Furthermore, archaeal uptake of glycerol from dead *Dunaliella* cells was documented in halite cores from the Dead Valley, California^63^ and the Saline Valley, California^64^. Although glycerol biosynthetic pathways were not found in the *Dolichomastix* genome from Salar Grande^15^, the algae may accumulate glycerol from the environment, released upon cell death when environmental conditions deteriorate.

In conclusion, one-year transplants showed that decreasing water availability was energetically more costly to the halite communities, particularly at the ALMA site, where communities experienced several cycles of hydration and dehydration. These extreme conditions favored opportunistic salt-in strategists equipped with mechanisms for energy conservation under high salt, such as the intracellular accumulation of KCl, the use of light-dependent proton pumps, and the ability to consume metabolites released from dead community members. Our work underscores how those fluctuations in water availability are important drivers of community structure and functional adaptations in the halite niche.

## METHODS

### Experimental design and sampling

Salar Grande (20.95 S, 70.02 W; 638 masl) was chosen as the “control site” for the transplant experiment. Halite microbial communities have been extensively characterized at this site and represented a suitable baseline^4,5,9,13,15^. In February 2018, halite nodules were transplanted from Salar Grande to the ALMA observatory on the Altiplano (23.07 S, 67.98 W; 2,885 masl), the hyper-arid Yungay region (24.05 S, 69.9 W; 953 masl), and the less arid Chañaral region in the southern part of the desert (26.17 S, 70.3 W; 848 masl) (Fig. 1). For each transplant, 8 nodules (3 nodules with environmental sensors and 5 nodules for metagenomic analyses) were collected within an area <100 m^2^. Eight nodules were marked in Salar Grande as controls. In March 2019, five transplanted nodules were harvested in the field at each site, and weather data were collected from all data loggers (see below). The nodules were cut in half with a circular saw, and the colonization zone was scrapped with a sterilized knife. The resulting grounded halite was collected in sterile Whirl-Pak bags and placed in dark and dry conditions for transportation and storage until analysis.

### Environmental data analysis

Atmospheric relative humidity (RH_air_) and temperature (T_air_) were recorded with HOBO Pro v2 External T/RH Data Loggers installed 1 meter above the ground at each of the four sites. Photosynthetic light was recorded at ground level at each site with a HOBO PAR (photosynthetic active radiation) sensor that measures light intensity in the range 0-2500 μmol.m^-2^.sec^-1^ over wavelengths from 400 to 700 nm. Halite temperature (T_halite_) and relative humidity (RH_halite_) were measured inside three transplanted nodules at each site using the same HOBO Pro v2 External T/RH Data Loggers. Sensors were installed at the top, middle, and bottom of the nodules by drilling three separate holes and sealing the sensor with resin as previously described^13^ (Fig. S1). All sensors were set to collect data every 30 minutes.

Data points for T, RH, and PAR were aligned to the same times on each recorded day by modeling raw data time-series and estimating values at 3-minute intervals, starting at midnight. Generalized Additive Models (GAMs) were built for T, RH, and PAR with adaptive smooths of the time of day expressed as day-hours in 3 minute intervals and maximum likelihood based optimization of the smoothness parameters^16^. Models were implemented with the GAM package^17^ in R (version 4.1.0). Daily total values for environmental variables were calculated independently for each sensor. Mean values between replicate halite nodules were reported for top, middle, and bottom positions.

The daily dynamics of T, RH, and PAR were used to calculate cumulative daily hours of wet and wet+light periods inside and outside halite nodules as proxies for potential metabolic activity. We assumed water was available to the halite communities when RH_halite_ ≥ 75%, the equilibrium relative humidity of a saturated NaCl brine^18^. For PAR, we used PAR ≥ 0.01 μmol photons m^-2^ sec^-1^ as the absolute minimum irradiance required by highly adapted primary producers^19^. We calculated the cumulative daily hours of wet and wet+light periods outside the nodules by assuming a RH_air_ threshold ≥ 95% and PAR ≥ 0.01 μmol photons m^-2^ sec^-1^. The RH_air_ threshold value is based on the observation that fog water collection in the Atacama Desert can occur at RH_air_ values slightly lower than 100%^20^ and to account for possible dew deposition^21^. For each day, the cumulative number of wet and wet+light hours were generated as a sum of 3-minute intervals at which values for RH and PAR exceeded respective thresholds.

### DNA extraction and sequencing

Genomic DNA (gDNA) was extracted from grounded halite using the DNeasy Powersoil DNA extraction kit (QIAGEN). The gDNA was cleaned with 3x AMPure XP Beads (Beckman Coulter) before library construction. The KAPA HyperPlus kit (Roche) was used to construct the genomic libraries with 10 ng of DNA, following the manufacturer’s instructions. Final library size selection was made with 0.5x and 0.7x KAPA HyperPure beads (Roche) ratio to recover fragments 300 bp −500 bp in length. Five libraries for Salar Grande and 3 for the other locations were paired-end sequenced by Novogene (https://en.novogene.com/) with 150 bp reads length using the Illumina NovaSeq 6000 platform.

### Sequences quality filtering, assembly, and binning

Sequence quality control, assembly, annotation, and quantification of contigs were performed using the MetaWRAP pipeline v1.3.2^22^. Raw reads were trimmed, and human contamination was removed using the read_qc module with default parameters. Two strategies for metagenome-assembled genomes (MAGs) assembly and binning were implemented (Fig. S2). The first strategy retrieved the less abundant MAGs as recommended in the metaWRAP workflow; reads were concatenated in a single file, assembled into contigs with the metaWRAP’s assembly module using MEGAHIT^23^, and binned into MAGs with the binning module using METABAT2^24^. The second strategy retrieved the most abundant MAGs; data from each sample were assembled individually to contigs using METASPADES^25^ and binned into MAGs using METABAT2. MAGs obtained from the two assembly strategies were pooled and de-replicated using dRep v2.0.0^26^ to get the most representative MAG for each species. Genomes were filtered by quality according to completeness (≥70%) and contamination (≤5%) using CheckM^27^. MAGs with an average nucleotide identity ≥ 95% and a coverage threshold ≥ 10% were considered the same species^28^.

### Taxonomic annotation and diversity analysis

Contigs and MAGs were quantified and normalized to sample size using the quant_bins module from metaWRAP. The resulting tables were transformed to relative abundance expressed as the percentage of each contig or bin in its sample using the vegan package in R^29^. Taxonomic annotation at the phylum level was performed with contigs and MAGs using the classify_bins module in metaWRAP and the NCBI nucleotide database v4.0. Phyla relative abundances were calculated for each site, ratios of archaea to bacteria were calculated by dividing relative abundances for each site, and the results were visualized by boxplots. Significance of differences for means of archaea: bacteria ratio and phyla relative abundances were evaluated using Welch’s t-test. The Bray-curtis index for each site was calculated using the vegan package. Distance matrices were visualized in PCoA plots generated with the ape R package^30^. The significance of clusters in PCoA plots was assessed with the ANOSIM statistic calculated in vegan.

### Functional annotation

Contigs obtained from the METASPADES assemblies were uploaded to the Integrated Microbial Genomes/Metagenomes from the Joint Genome Institute (JGI IMG) v5.0.0 service^31^. Gene copy counts were downloaded from the server with their assigned KO terms list and linked to their respective KEGG BRITE pathway classification. Counts were normalized by dividing by the number of reads in the corresponding sample and transformed to relative abundance using vegan. The 40 most abundant pathway relative abundances were visualized in a heatmap using the pheatmap R package. The average isoelectric point (pI) was calculated for each metagenomic assembly; open reading frames (ORFs) were predicted with PRODIGAL^32^ using the annotate_bins module from metaWRAP, and the pI for each ORF was calculated using ProPAS^33^. For potassium transport potential (*tr*K genes), relative abundance of contigs annotated with the KOs: K03498 and K03499 were retrieved from the annotation table for each sample and visualized by boxplots. Significant differences among locations were assessed with Welch’s t-test for the pI and trK relative abundances.

### Specialized MAGs characterization

Specialized MAGs were defined as those with a mean relative abundance higher than 10% in at least one site; they were selected to be classified at a deeper taxonomic level (Table S3). Selected genomes were uploaded to GTDB-tk server^34^ to be taxonomic classified with the Genome Database Taxonomy (GTDB)^35^. Closely related MAGs at the family level were retrieved from the GTDB for phylogenomic tree construction using GTOTree v1.6.11^36^ with MUSCLE alignment of single-copy genes (SCP) in the HMM database and FastTree algorithm for tree construction.

Potential osmoadaptation strategies for specialized MAGs were obtained by uploading protein prediction from annotate_bins in metaWRAP to the GhostKOALA server and retrieving the list of KO identifiers for each MAG^37^. KO lists were linked to (1) osmoadaptation KO genes lists based on gene lists from Wong, et al. ^38^ and (2) the utilization of glycerol as a carbon source based on gene lists from Wong, et al. ^38^ and Oren ^39^, respectively. Occurrences of osmoadaptation strategies and use of glycerol were represented in a heatmap indicating the presence or absence of each gene in the selected MAGs.

### Algal and viral contigs quantification

The *Dolichomastix* alga genome, previously assembled by Uritskiy, et al. ^15^, was retrieved from github (https://github.com/ursky/metatranscriptome_paper/tree/master/MAGS). Its representation in each sample, as genome copies per million of reads, was calculated using the quant_bins module from MetaWRAP and transformed to relative abundance using vegan in R; significance of variation across sites was calculated using Welch’s t-test.

Halite communities assembled contigs obtained from METASPADES assemblies were used for viral identification of dsDNA viruses, *Nucleocytoviricota* (NCLDV), and *Lavidaviridae* viruses using virsorter2^40^. Assembled contigs were quality checked with CheckV^41^ and taxonomically annotated with Cenote-Taker2 using default parameters^42^. Viral contig relative abundances were calculated by linking the contig annotation from Cenote-Taker2 with the contig depth obtained with metaWRAP quant_bins module. Variation of virus relative abundance across locations was represented in boxplots, and the significance was assessed using Welch’s t-test.

## ACKNOWLEDGEMENTS

This work was funded by NASA grants 80NSSC19K0470 and NNX15AP18G. We thank the personnel at ALMA for their fieldwork support, hospitality, and access to the facility. We thank Peter R. McCullough and Dana Burton for field work assistance. McCullough’s travel was supported in part by the Space Telescope Science Institute’s Director’s Discretionary Fund under grant D0001.82413.

## AUTHOR CONTRIBUTIONS

JD and AD conceptualized and designed the study; JD, AD, CP, and PW collected in-field samples; CP processed and sequenced samples; CP and JD analyzed the molecular data; AD and PW analyzed the climate data. All wrote the manuscript.

## DATA AVAILABILITY

Raw sequences are available at the National Center for Biotechnology Information (NCBI) under the BioProject ID PRJNA808683. BioSample accession numbers are: SAMN26101366 - SAMN26101368 for ALMA, SAMN26101369 - SAMN26101371 for Chañaral, SAMN26101372 - SAMN26101376 for Salar Grande, and SAMN26101377 - SAMN26101379 for Yungay. All analysis pipelines, processed data, analysis and visualization scripts, and reconstructed MAGs are available at https://github.com/capfz200/Halite_paper. The metagenome assembly and functional annotation are available in the Joint Genome Institute (JGI) Integrated Microbial Genomes and Microbiomes portal under the GOLD Study ID Gs0154137.

