## Supplementary material for "Adaptations of endolithic communities to abrupt environmental changes in a hyper-arid desert": FigS1

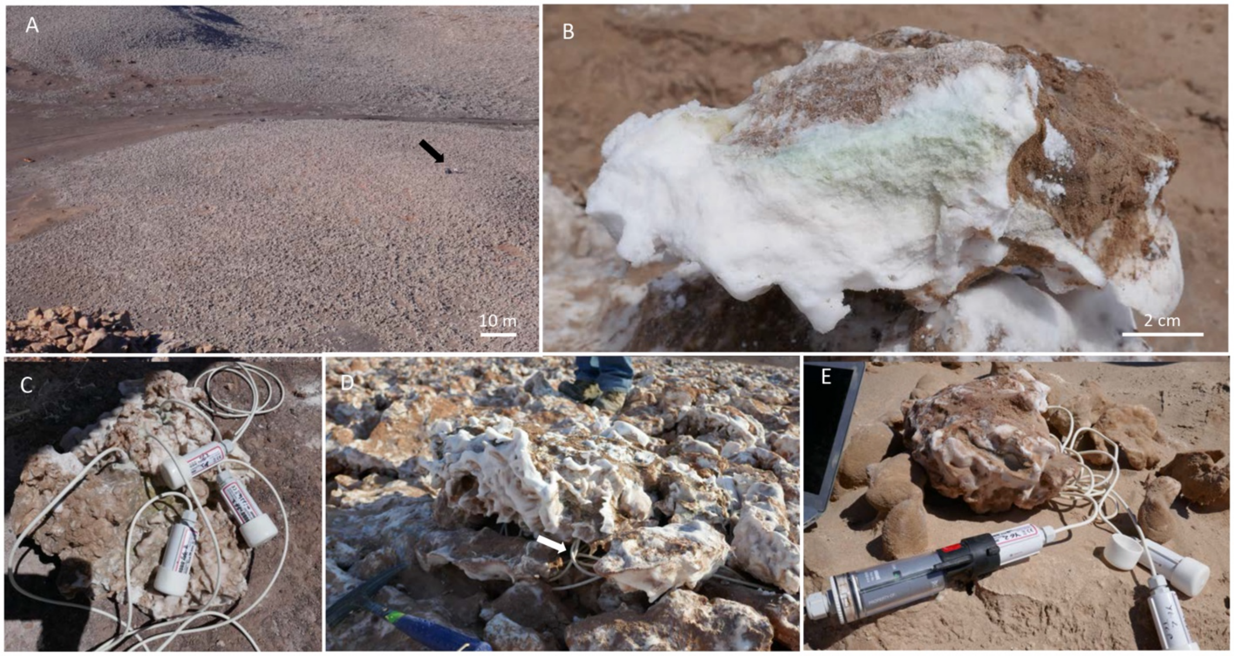


Figure S1. Field photos. (A) Salar Grande landscape with field site (black arrow). (B) Cross-section of a halite nodule showing the green colonization zone. (C) Bottom view of a halite module with internal sensors sealed in the salt, wires, and Hobo recorders. (D) Sensor-equipped halite nodule in the field with apparent wires (white arrow). (E) Data downloading from internal sensors in Yungay.

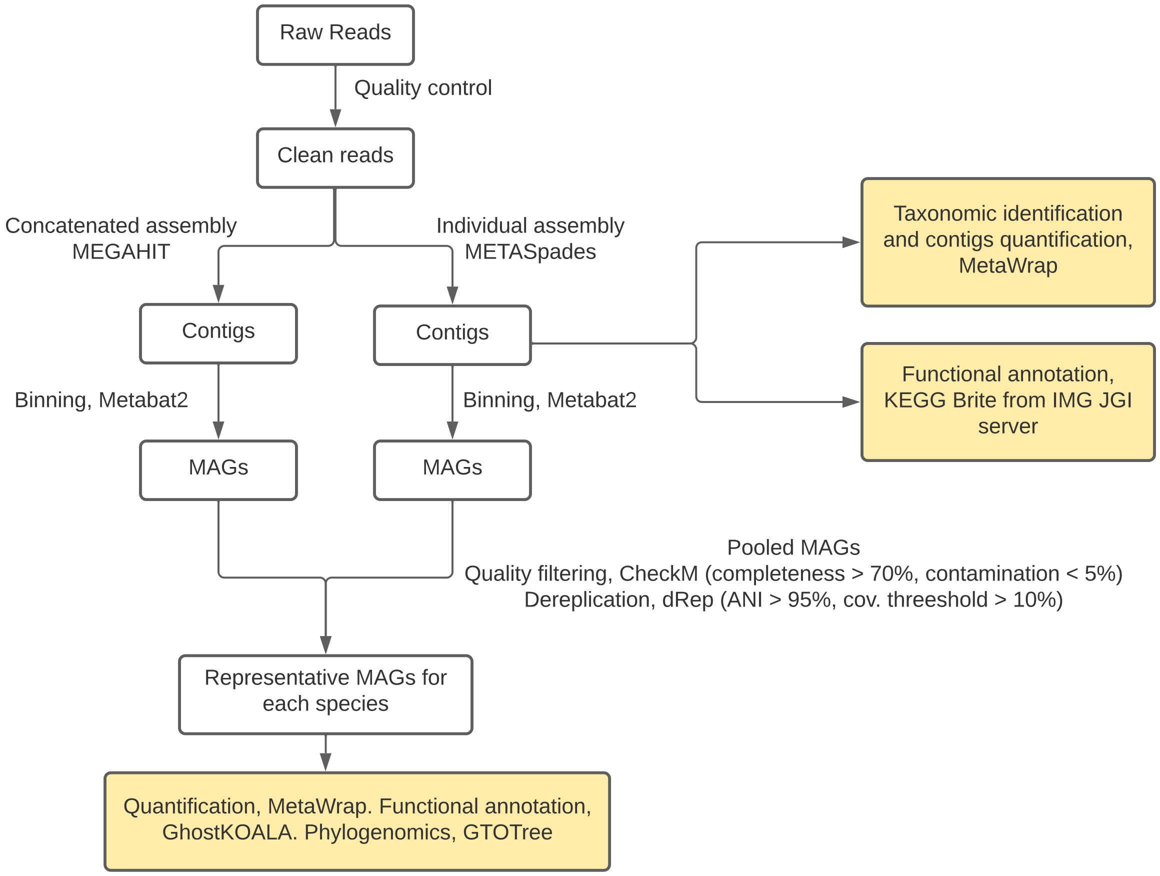


Figure S2. Workflow of the metagenomic analysis.

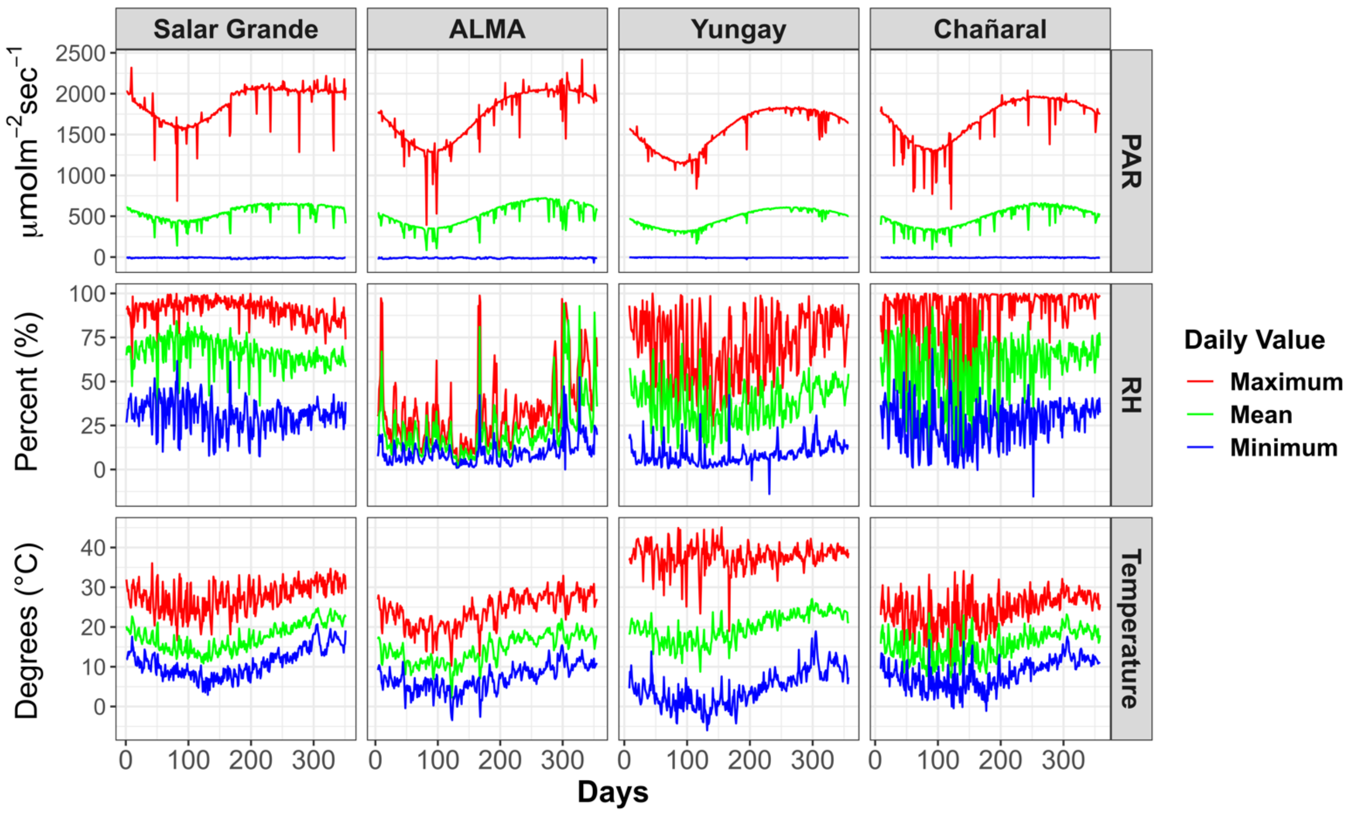


Figure S3. Environmental atmospheric conditions (outside halite nodules) at different sites.

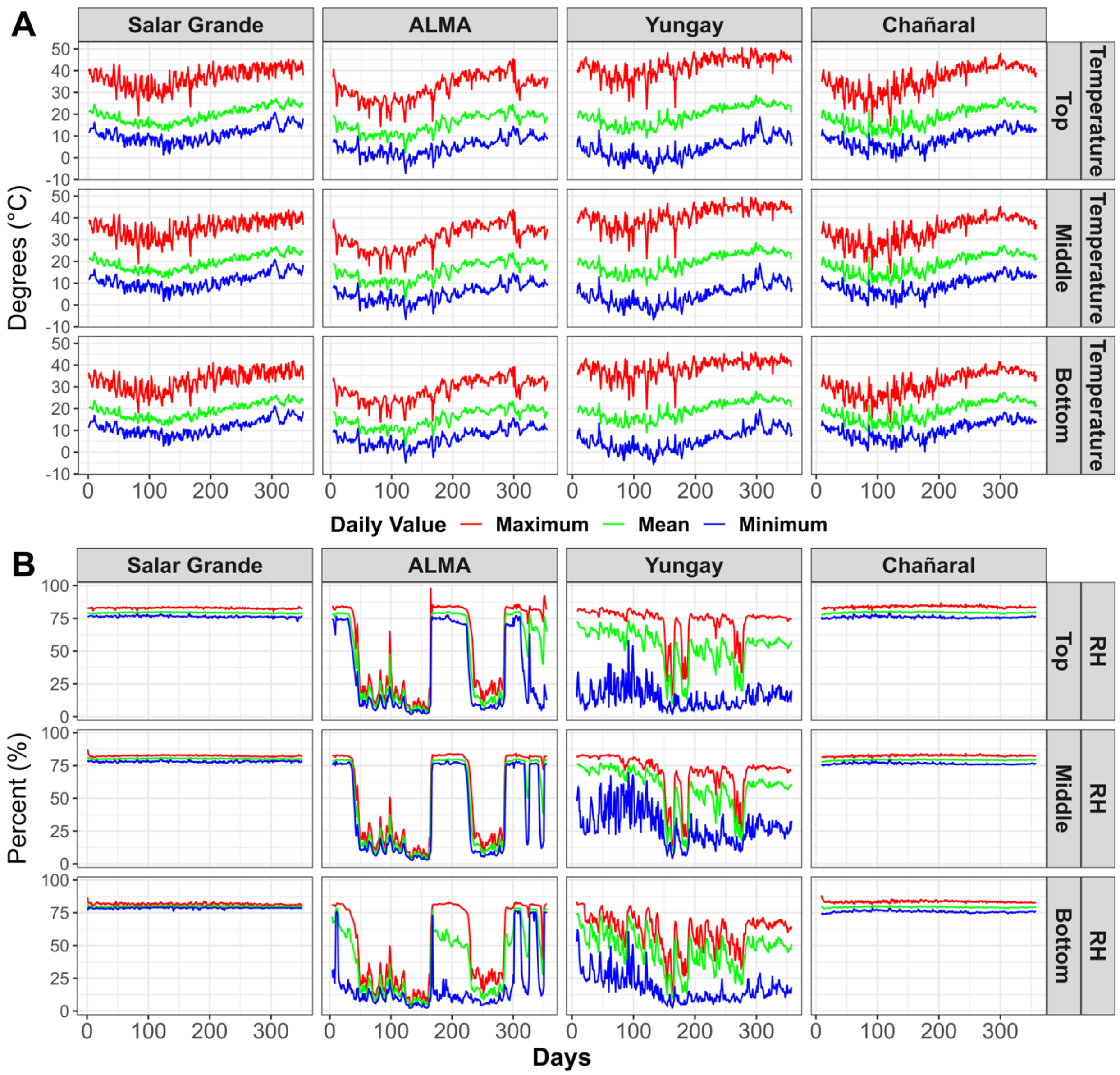


Figure S4. Environmental conditions inside halite nodules at different sites.

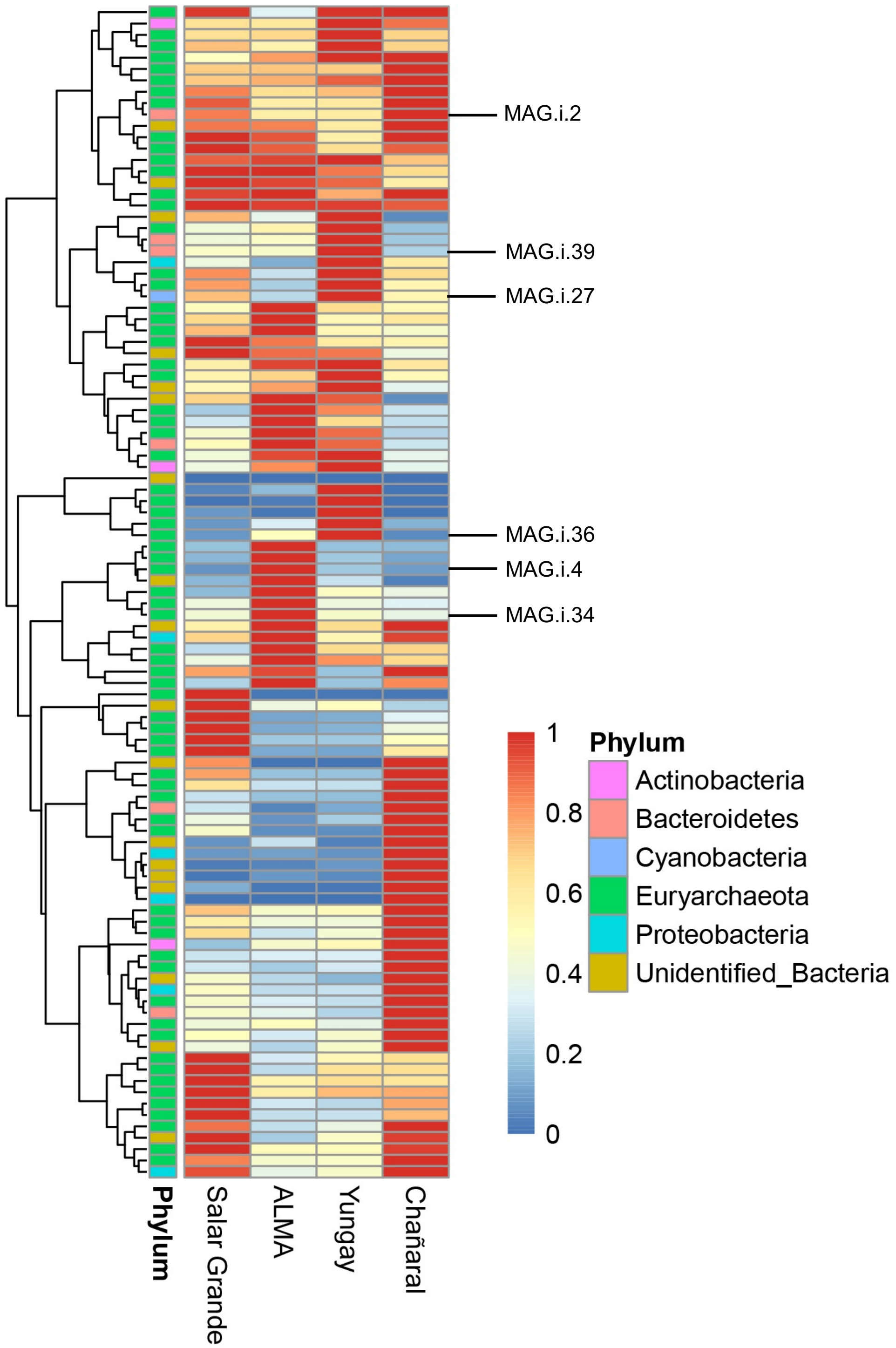


Figure S5. Taxonomic annotation and relative abundance of MAGs across sites standardized to the maximum value in each row. Annotated are the specialized MAGs.

A

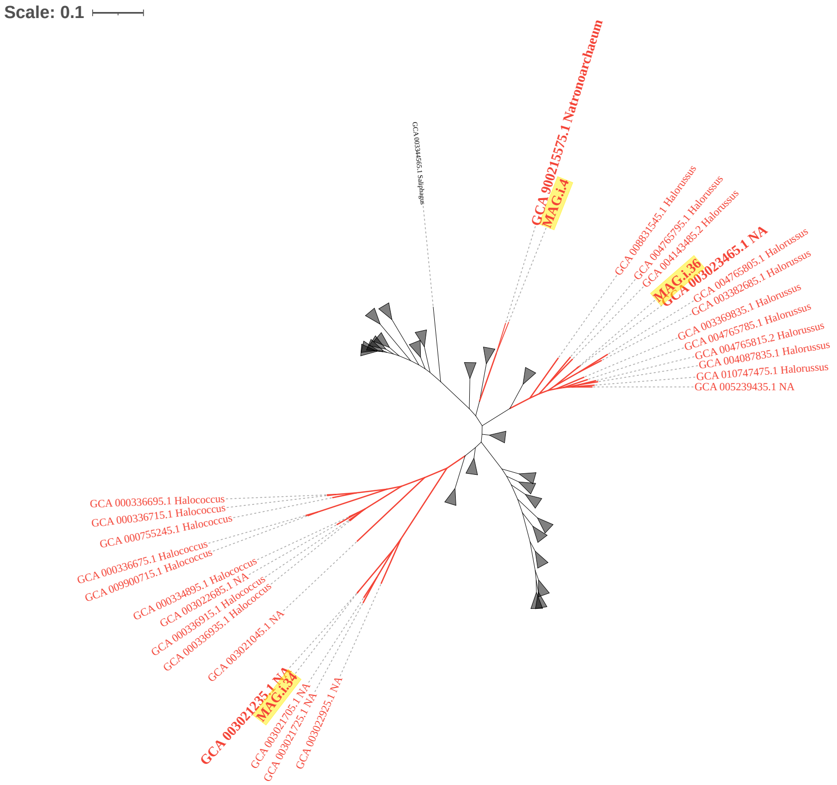

Scale: 0.1
GCA 900215575.1 Natronoarchaeum
MAG.i.4
GCA 003023465.1 NA
MAG.i.36
GCA 004765805.1 Halorussus
GCA 003382685.1 Halorussus
GCA 003369835.1 Halorussus
GCA 004765785.1 Halorussus
GCA 004765815.2 Halorussus
GCA 004087835.1 Halorussus
GCA 010747475.1 Halorussus
GCA 005239435.1 NA
GCA 000336695.1 Halococcus
GCA 000336715.1 Halococcus
GCA 000755245.1 Halococcus
GCA 000336675.1 Halococcus
GCA 009900715.1 Halococcus
GCA 000334895.1 Halococcus
GCA 003022685.1 NA
GCA 000336915.1 Halococcus
GCA 000336935.1 Halococcus
GCA 003021045.1 NA
GCA 003021235.1 NA
MAG.i.34
GCA 003021705.1 NA
GCA 003021725.1 NA
GCA 003022925.1 NA


B

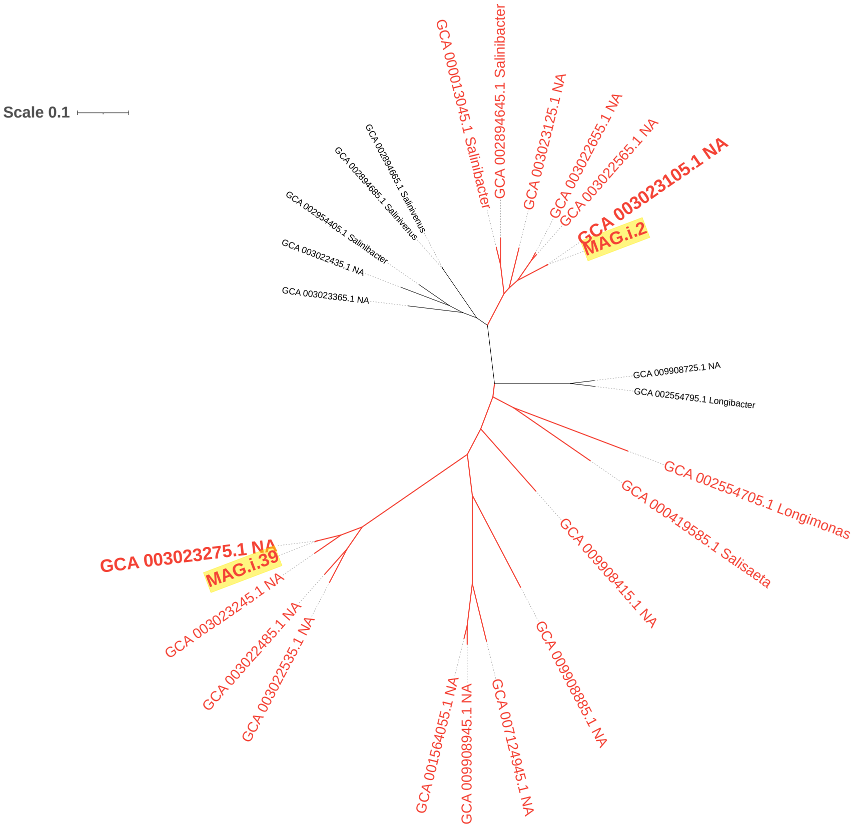

Scale 0.1
GCA 000013045.1 Salinibacter
GCA 002894645.1 Salinibacter
GCA 003023125.1 NA
GCA 003022655.1 NA
GCA 003022565.1 NA
GCA 003023105.1 NA
MAG.i.2
GCA 002894605.1 Salinivenus
GCA 002894685.1 Salinivenus
GCA 002954405.1 Salinibacter
GCA 003022435.1 NA
GCA 003023365.1 NA
GCA 009908725.1 NA
GCA 002554795.1 Longibacter
GCA 002554705.1 Longimonas
GCA 000419585.1 Salisaeta
GCA 009908415.1 NA
GCA 009908885.1 NA
GCA 007124945.1 NA
GCA 009908945.1 NA
GCA 001564055.1 NA
GCA 003022535.1 NA
GCA 003022485.1 NA
GCA 003023245.1 NA
GCA 003023275.1 NA
MAG.i.39

C

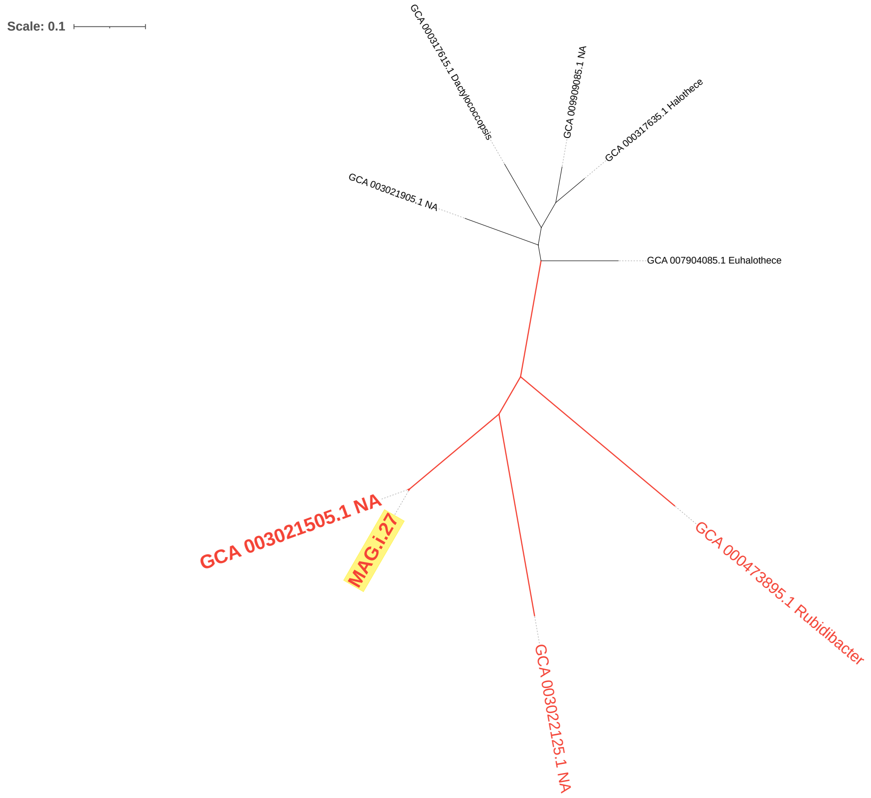


Figure S6. Phylogenomic trees for (A) *Halobacteriales*, (B) *Salinibacteraceae*, and (C) *Rubidibacteraceae*. Scale is the number of substitutions per site.

Table S1. Average Nucleotide Identity (ANI) of specialized MAGs and Genome Database Taxonomy (GTDB)

| MAG | FastANI Taxonomy | FastANI ANI (%) | Fraction aligned | Reference |
|---|---|---|---|---|
| MAG.i.39 | d_Bacteria;p_Bacteroidota;c_Rhodothermia;o_Rhodothermales;f_Salinibacteraceae | 98.32 | 0.93 | GCA_003023275.1 |
| MAG.i.2 | d_Bacteria;p_Bacteroidota;c_Rhodothermia;o_Rhodothermales;f_Salinibacteraceae; g_Salinibacter | 99.03 | 0.82 | GCA_003023105.1 |
| MAG.i.36 | d_Archaea;p_Halobacterota;c_Halobacteria;o_Halobacteriales;f_Haloadaptaceae | 98.46 | 0.94 | GCA_003023465.1 |
| MAG.i.34 | d_Archaea;p_Halobacterota;c_Halobacteria;o_Halobacteriales;f_Halococcaceae | 98.71 | 0.91 | GCA_003021235.1 |
| MAG.i.4 | d_Archaea;p_Halobacterota;c_Halobacteria;o_Halobacteriales;f_Natronoarchaeaceae; g__Natronoarchaeum | <95%* | na | GCF_900215575.1 |
| MAG.i.27 | d_Bacteria;p_Cyanobacteria;c_Cyanobacteriia;o_Cyanobacteriales;f_Rubidibacteraceae | 99.16 | 0.95 | GCA_003021505.1 |

* MAG.i.4 taxonomy was confirmed using the relative evolutionary divergence method; amino acid identity was 95.75% and RED value was 0.92; na, non-applicable
